# A single 2’-O-methylation of ribosomal RNA gates assembly of a functional ribosome

**DOI:** 10.1101/2022.02.16.480697

**Authors:** James N. Yelland, Jack P.K. Bravo, Joshua J. Black, David W. Taylor, Arlen W. Johnson

## Abstract

RNA modifications are widespread in biology, and particularly abundant in ribosomal RNA. However, the significance of these modifications is not well understood. We show that methylation of a single universally conserved nucleotide, in the catalytic center of the large subunit, gates ribosome assembly. Massively parallel mutational scanning of the essential nuclear GTPase Nog2 identified important interactions with ribosomal RNA helix 92, particularly with the methylated A-site base Gm2922. We found that 2’-O-methylation of G2922 is needed for efficient nuclear export of the large subunit. Critically, we identified single amino acid changes in Nog2 that completely bypass its dependence on G2922 methylation. By solving the cryo-EM structure of the unmodified nascent subunit, we reveal how methylation flips Gm2922 into the active site channel of Nog2. This work demonstrates that a single RNA modification is a critical checkpoint in ribosome biogenesis, and suggests that RNA modifications can play an important role in regulation and assembly of macromolecular machines.

## Main

Chemical modification of RNA is pervasive in all three domains of life and is abundant in ribosomal RNA (rRNA). Most rRNA modifications cluster around functionally important regions of the ribosome, including the decoding center and the peptidyl transferase center (PTC), where they are thought to stabilize folding and tertiary structure of RNA at these functionally important sites^1–3^. A noteworthy example is the universally conserved A-site loop of rRNA helix 92 (H92), which is highly modified and contains the 2’-O-methylated guanosine (Gm2922 in yeast). During translation, Gm2922 base-pairs with the 3’ end of an accommodated tRNA to position it in the A-site for the peptidyl transferase reaction^4^. However, the importance of methylating G2922, and indeed of any specific rRNA modification, remains poorly understood.

Eukaryotic ribosome assembly is a complex and intricately organized pathway spanning multiple cellular compartments^5,6^. Over 200 assembly factors play a role in assembling the functional ribosome, including several evolutionarily conserved ATPases and GTPases which drive rRNA rearrangements and progression of ribosome assembly. One such enzyme is Nog2, an essential nuclear GTPase that is conserved between yeast and humans. Nog2 is required during critical RNA rearrangements within the precursor large (pre-60S) subunit, prior to nuclear export of the subunit^7^. However, the specific function of Nog2 has not been identified. To better understand the function of this GTPase, we searched for functionally important regions of yeast Nog2 using massively parallel mutagenic scanning. Unexpectedly, we discovered that physical interaction of Nog2 with H92, including recognition of the 2’-O-methylated base Gm2922, is critical for cell viability. We found that pre-60S subunits lacking methylated G2922 are blocked for assembly and nuclear export, and that orthogonal 2’-O-methylation of G2922 or single amino acid changes in Nog2 overcame the ribosome assembly and nuclear export defects of pre-60S lacking methylated G2922. Finally, we used cryo-electron microscopy to solve the structure of a nascent 60S subunit from cells with unmethylated G2922, showing that an unmethylated G2922 fails to engage with Nog2. This work thus demonstrates an important role for a single rRNA modification as a structural checkpoint in the eukaryotic ribosome assembly pathway.

### Physical interaction between H92 and Nog2 is important for viability in yeast

To identify functionally important regions of Nog2, we generated a library of ∼7,800 *NOG2* codon variants (∼26% of possible codon substitutions) via mutagenic PCR. We introduced this library into a yeast strain in which we could select for functional mutants, where the endogenous *NOG2* gene was under control of the glucose-repressible *GAL1* promoter. We sequenced and analyzed *NOG2* variants with a deep mutagenesis scanning pipeline^8^ (Figure S1A-S1D) and mapped intolerance to mutation onto the structure of Nog2 (Figure 1A). Surprisingly, we found a distinct resistance to mutation in the amino acid residues interacting with H92 (Figure 1B). More strikingly, several mutation-intolerant residues surround the 2’-O-methylated guanosine base Gm2922 (Figure 1C). Published structures of Nog2-bound pre-60S intermediates show that Gm2922 is flipped from its canonical position^9^ by a 2’-*endo* pucker, into a channel gating the active site of Nog2 (Figures 1C-D, Figure S2A). Modeling the 2’-O-methyl of Gm2922, which is clearly resolved in a previous cryo-EM structure of Nog2^10^, reveals how it is accommodated at the entrance to the active site (Figure 1D, Figure S2B). Additionally, arginine 389 of Nog2 (R389), which resisted amino acid substitutions (Figure 1B), stabilizes Gm2922 through a cation-π interaction with the guanosine base (Figure 1C, Figure S2C)^11,12^. On the opposite side of the channel, serine 208 (S208) forms a hydrogen bond with the 5’ phosphate of Ψ2923 (Figure 1C, Figure S2C). As expected, S208 was also resistant to mutation in our analysis (Figure 1B). To directly assess the functional significance of Nog2 binding to Gm2922, we tested the ability of the strongly disfavored *NOG2* variants S208A and R389S to complement loss of endogenous *NOG2* (Figure 1E). Although we observed no obvious growth defect in the S208A variant, and a modest growth reduction in the R389S variant, combining these mutations rendered a lethal variant. These results demonstrate that at least two amino acids physically interacting with Gm2922 are interdependent and important for the essential function of Nog2.

**Figure 1.**
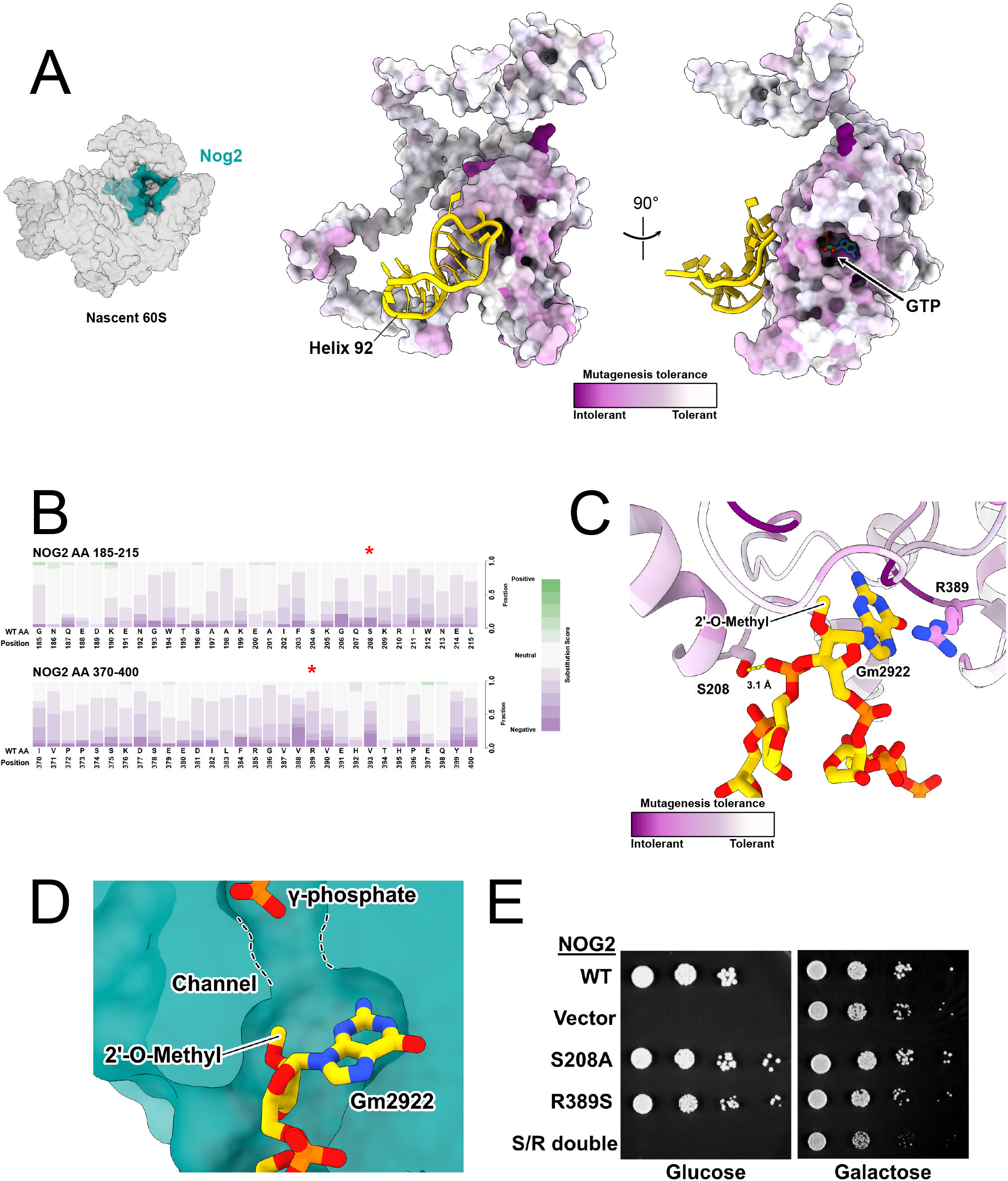
Essential function of Nog2 depends on physical interaction with H92. **A**. (Left) Structure of Nog2 relative to pre-60S (PDB 3JCT) in “crown view”. (Right) Nog2 colored by per-residue average score of amino acid tolerance to mutation. H92 is colored gold. **B**. Cumulative per-residue fitness scores for all amino acid substitutions at H92-interacting regions of Nog2. Residues S208 and R389 are denoted by red asterisks. **C**. Cartoon representation of Nog2 residues interacting with H92, colored as in (A). **D**. Cutaway surface representation of Nog2, with Gm2922 flipped into the active-site channel. The γ-phosphate of GTP is also shown. **E**. Serial dilution assays to test complementation by the indicated *NOG2* alleles in a repressible *pGAL1:NOG2* strain (glucose, endogenous *NOG2* repressed; galactose, expressed).

### Gm2922 is critical for ribosome assembly and nuclear export

G2922 is 2’-O-methylated by the conserved methyltransferase Spb1^13,14^, which is essential in yeast^15^. Previous studies showed that a catalytically dead variant of Spb1 is detrimental to cell growth^13,14^, but did not address the role of its target base. To specifically test the importance of methylating G2922, we engineered an artificial snoRNA to guide methylation of G2922 (Figure 2A). This snoRNA, designated *SNRG2922*, was designed to hybridize with nascent 25S rRNA and direct 2’-O-methylation of G2922 via the box C/D snoRNP complex^16^. We introduced *SNRG2922* into a yeast strain expressing a methyltransferase-deficient mutant of *SPB1, spb1-D52A*, which cannot methylate G2922^13,14^. Strikingly, expression of *SNRG2922* supported near wild-type growth in the *spb1-D52A* strain (Figure 2B). To verify that improved growth correlated with restoration of ribosome assembly, we analyzed the polysome profiles of wild-type *SPB1, spb1-D52A* and the *SNRG2922*-complemented *spb1-D52A* strains. The strong 60S synthesis defect of the *spb1-D52A* mutant, indicated by low levels of free 60S and polysomes, was greatly alleviated by expressing *SNRG2922* (Figure 2C). To confirm that *SNRG2922* directs methylation of Gm2922, we designed a primer extension assay, taking advantage of the fact that 2’-O-methylated rRNA is resistant to cleavage by RNAse T1^14^ (Figure S3A). G2922 was susceptible to cleavage in the *spb1-D52A* strain, but resistant to cleavage when *SNRG2922* was expressed, indicating that *SNRG2922* restores 2’-O-methylation of Gm2922 (Figure 2D). Together, these results demonstrate that 2’-O-methylation of G2922 is critical for ribosome biogenesis in yeast.

**Figure 2.**
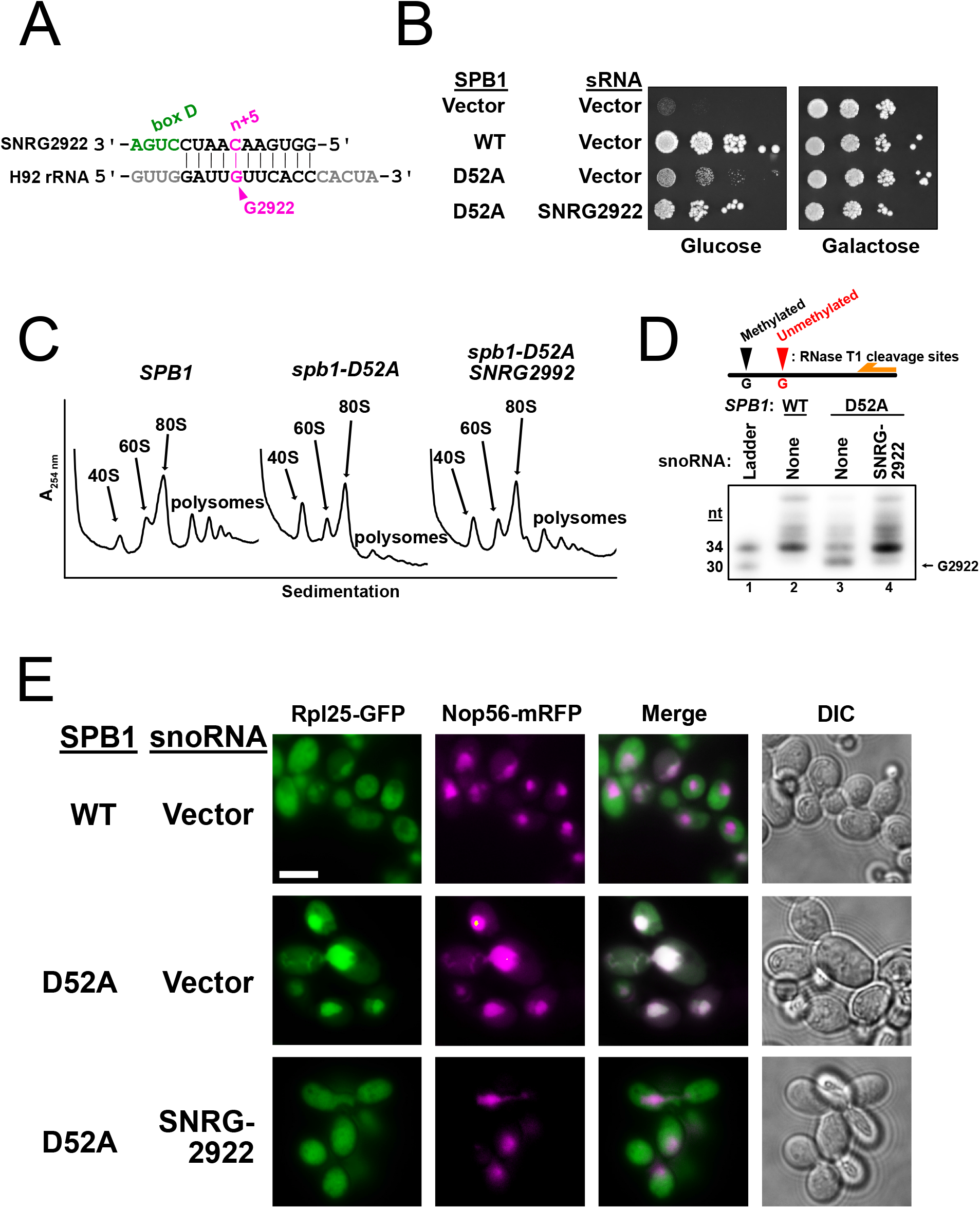
2’-O-methylation of Gm2922 is necessary and sufficient for robust cell growth and nuclear export of the nascent large ribosomal subunit. **A**. Design of synthetic snoRNA gene *SNRG222* to direct 2’-O-methylation of Gm2922, catalyzed by the box C/D snoRNP complex at the fifth nucleotide upstream of box D. **B**. Expression of *SNRG2922* overcomes the growth defect of the methyltransferase-defective *spb1-D52A* mutant in a *pGAL1:SPB1* strain (glucose, endogenous *SPB1* repressed; galactose, expressed). **C**. Sucrose gradient sedimentation profiles for the indicated *SPB1* and snoRNA alleles. **D**. Primer extension assay to probe 2’-O-methylation at G2922. Autoradiography of PAGE-separated primer extension products of RNAse T1-digested RNA. **E**. Fluorescence microscopy to monitor localization of large subunit (Rpl25-eGFP) and nucleolus (Nop56-mRFP). Scale bar = 5 µm.

Because Nog2 is required for export of the pre-60S^15,17^, we hypothesized that failure of G2922 to engage with Nog2 would interrupt nuclear export. To test this hypothesis, we monitored the intracellular localization of Rpl25-GFP, an established reporter for nuclear export of the pre-60S^18^. While Rpl25 was predominantly cytoplasmic in cells expressing wild-type *SPB1*, reflecting the localization of mature ribosomes, the methyltransferase-deficient *spb1-D52A* induced nucleolar localization of Rpl25-GFP, suggesting that lack of 2’-O-methylation leads to nucleolar retention of the pre-60S (Figure 2E). Remarkably, expression of *SNRG2922* restored localization of Rpl25-GFP to the cytoplasm (Figure 2E), indicating that the single 2’-O-methlylation of Gm2922 is required for efficient nuclear export of the pre-60S.

### Single amino acid changes in Nog2 bypass dependence on Gm2922

We hypothesized that if Nog2 associates weakly with an unmodified G2922, overexpression of Nog2 would overcome cellular dependence on Gm2922. Indeed, overexpression of Nog2 weakly suppressed the growth defect of methyltransferase-deficient *spb1-D52A* cells (Figure 3A). Similarly, we reasoned that variants of Nog2 capable of binding more strongly to H92 would more efficiently bypass the growth defect caused by lack of G2922 methylation. To directly test this, we designed a *NOG2* variant, threonine 195 to arginine (T195R), a positively charged substitution predicted to bind more strongly to the negatively charged H92 phosphodiester backbone (Figure 3B, left). Strikingly, *NOG2-T195R* restored growth in the *spb1-D52A* strain to near wild-type levels (Figure 3A). In parallel, we screened our *NOG2* variant library for randomly-generated suppressors of *spb1-D52A*, and identified a second *NOG2* variant, histidine 392 to arginine (H392R), which also restored growth to near wild-type levels (Figure 3A). As predicted, this variant should also increase electrostatic binding with the phosphodiester backbone of H92 (Figure 3B, right). We confirmed that expression of *NOG2-H392R* restored 60S biogenesis in the *spb1-D52A* strain (Figure 3C), despite the absence of 2’-O-methylated G2922 (Figure 3D). Moreover, expression of *NOG2-H392R* restored nuclear export of the pre-60S (Figure 3E). Our finding that single amino acid changes in Nog2 can bypass cellular dependence on Gm2922 strongly suggests that Nog2 directly monitors formation of Gm2922 to gate nuclear export of the large ribosomal subunit.

**Figure 3.**
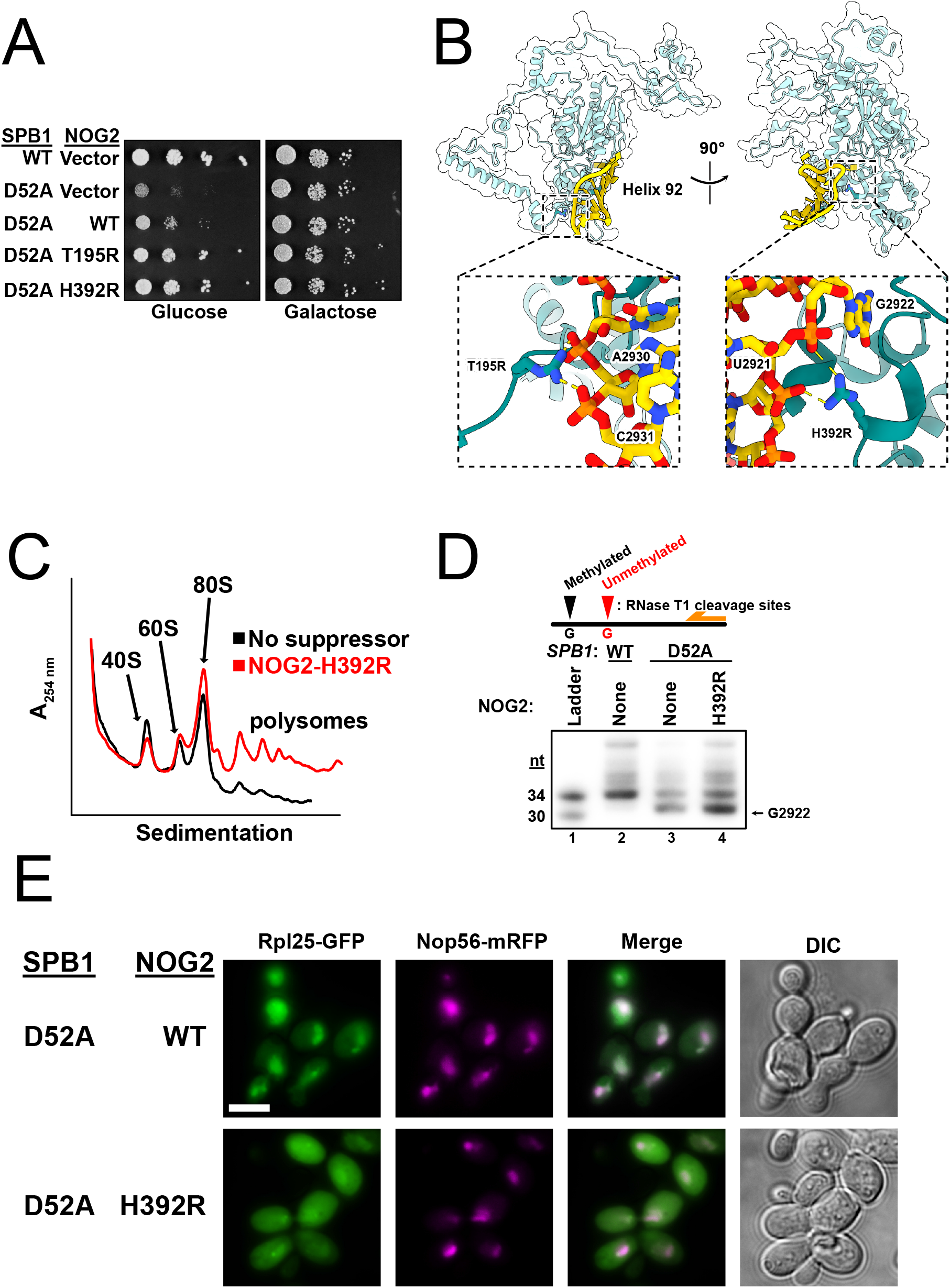
Single amino acid changes in Nog2 restore robust growth and nuclear export in the absence of Gm2922. **A**. Genetic suppression of a lack of Gm2922 by the indicated *NOG2* alleles in a *pGAL1:SPB1* strain (glucose, endogenous SPB12 repressed; galactose, expressed). **B**. Cartoon and surface representation of Nog2, with suppressing amino acid changes modeled (PDB 3JCT). Left inset: Suppressing amino acid change T195R modeled to show novel electrostatic interactions with the phosphates of H92 bases A2930 and C2931. Right inset: Suppressing amino acid change H392R modeled similarly to show novel electrostatic interactions with the phosphates of H92 bases U2921 and G2922. **C**. Sucrose gradient sedimentation profile from *spb1-D52A* cells suppressed by *NOG2-H392R*, contrasted with empty vector (black trace same as shown in Figure 2C). **D**. Primer extension assay to probe 2’-O-methylation at G2922 in WT cells, cells expressing methyltransferase-defective *spb1-D52A*, and *spb1-D52A* suppressed by *NOG2-H392R*. Lanes 2-3 are the same samples as loaded in Figure 2D. **E**. Fluorescence microscopy to monitor location of large subunit (Rpl25-eGFP) and nucleolus (Nop56-mRFP) when spb1-D52A is suppressed by WT *NOG2* or *NOG2-H392R*. Scale bar = 5 µm.

### Unmethylated G2922 does not engage with Nog2

To investigate the molecular defects caused by lack of G2922 modification, we genetically tagged Nog2 and immunopurified Nog2-bound pre-60S subunits from cells expressing methyltransferase-deficient *spb1-D52A*, or wild-type *SPB1* as a control (Figure S4A). Semi-quantitative tandem mass-spectrometric analysis showed that Nog2 particles from *spb1-D52A* cells are depleted of components of the Rix1-Rea1 complex required for maturation of the 60S central protuberance^19^, as well as of Sda1, a ribosome assembly factor involved in recruitment of the Rix1-Rea1^12^ complex, and nuclear export of the pre-60S subunit^20^ (Figure S4B). Furthermore, we observed that unmodified subunits are deficient in Arx1 and Bud20, nonessential but important assembly factors that bind the nuclear subunit and contribute to its efficient cytoplasmic export^21–23^ (Figure S4A, Figure S4B). In turn, Nog2 particles from *spb1-D52A* cells are enriched for nucleolar factors, including Spb1, consistent with arrest of an earlier pre-ribosomal complex (Figure S4B). Failure to methylate G2922 therefore traps a nucleolar intermediate, supporting our conclusion that Gm2922 is important for the pre-60S to transition from the nucleolus to the nucleoplasm.

Our results strongly suggest that Nog2 binding to H92 depends on methylation of G2922. To directly visualize the consequences of unmethylated G2922, we used cryo-electron microscopy (cryo-EM) to solve the structure of an unmodified pre-60S subunit bound by Nog2 at an overall resolution of 2.9 Å (Figure 4A, Figure S5 and Figure S6, Table 1). In our structure, density for Nog2 largely agrees with prior structure determinations^10,12,19^ with switch I and switch II (motifs G2 and G3) ordered around GTP clearly present in the active site (Figure S6A). Importantly, however, the unmodified G2922 is not flipped into the active site channel of Nog2, unlike the modified Gm2922 in previously solved structures (Figure 4B-C). Instead, when G2922 lacks a 2’-O-methyl group, H92 closely matches the conformation it adopts within the mature large ribosomal subunit, where Gm2922 stacks on Ψ2923 (Figure S6B)^9^. This result unambiguously shows that methylation of G2922 is critical for its engagement with the active site channel of Nog2. Thus, Nog2 directly assesses the methylation status of Gm2922, gating ribosome biogenesis prior to nuclear export of the nascent 60S subunit.

**Table 1.** Cryo-EM data collection, refinement and validation statistics.

|  |  |
| --- | --- |
|  | #1 Nog2-3xFLAG,<br>unmodified G2922<br>(EMDB-xxxx)<br>(PDB xxxx) |
| **Data collection and processing** |  |
| Magnification | 29,000 x |
| Voltage (kV) | 300 |
| Electron exposure (e-/Å<sup>2</sup>) | 70 |
| Defocus range (μm) | -1.5 - -2.5 |
| Pixel size (Å) | 0.81 |
| Symmetry imposed | C1 |
| Initial particle images (no.) | 403,566 |
| Final particle images (no.) | 15,954 |
| Map resolution (Å) | 3.1 |
| FSC threshold | 0.143 |
| Map resolution range (Å) | 2 - 10 |
| **Refinement** |  |
| Initial model used (PDB code) | 3JCT |
| Model resolution (Å) | 2.9 |
| FSC threshold | 0.5 |
| Model resolution range (Å) | 2.5 |
| Map sharpening *B* factor (Å<sup>2</sup>) | 41.4 |
| Model composition |  |
| Non-hydrogen atoms | 140,964 |
| Protein residues | 8,739 |
| Nucleotides | 3,337 |
| *B* factors (Å<sup>2</sup>) |  |
| Protein | 70.19 |
| Nucleotide | 85.64 |
| R.m.s. deviations |  |
| Bond lengths (Å) | 0.003 |
| Bond angles (°) | 0.626 |
| Validation |  |
| MolProbity score | 2.11 |
| Clashscore | 13.48 |
| Poor rotamers (%) | 2.98 |
| Ramachandran plot |  |
| Favored (%) | 97.48 |
| Allowed (%) | 2.50 |
| Disallowed (%) | 0.02 |

**Figure 4.**
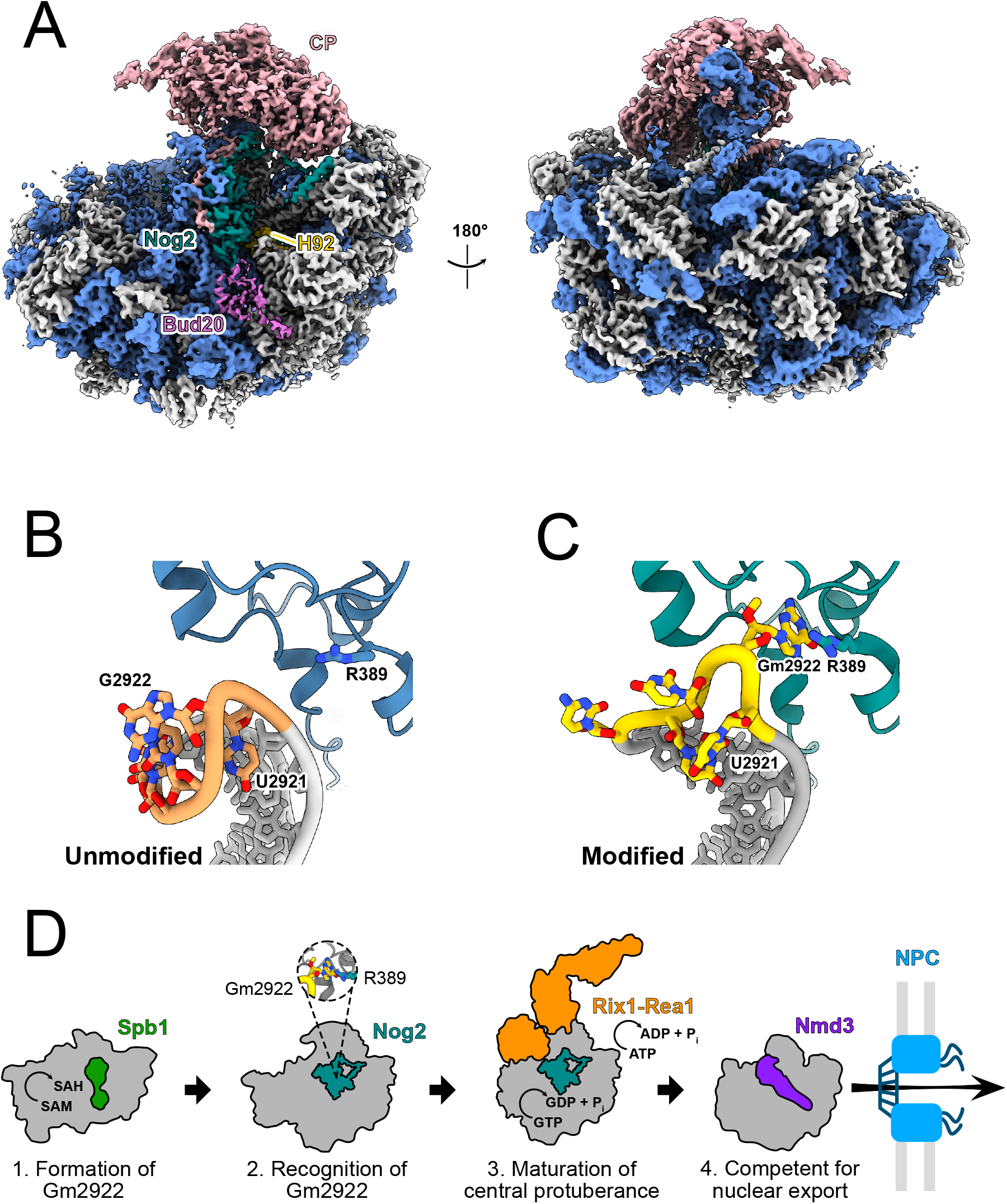
Nog2 does not engage with unmodified G2922. **A**. Cryo-EM structure of pre-60S in complex with Nog2 bound to H92 from methyltransferase-defective *spb1-D52A* mutant cells in which G2922 is unmodified (crown view, left). The unrotated central protuberance (CP) is indicated along with Nog2, H92 and Bud20. **B-C**. Atomic models of unmodified G2922 and Nog2 **(B)**, and modified G2922 and Nog2 (PDB 3JCT) **(C). D**. Cartoon model for Nog2 gating of 60S assembly and nuclear export, dependent on methylation of G2922.

## Discussion

Our work demonstrates how a single nucleotide modification serves as a structural checkpoint in ribosome assembly. We have shown that recognition of Gm2922 by the essential GTPase Nog2 couples structural maturation of the ribosome to Nog2-dependent nuclear export. In this scenario, methylation of G2922 by Spb1 initially assesses the correct folding of the RNA of the proto A-site. Nog2 “reads” the modification status of Gm2922 to ensure that only ribosomes with a methylated G2922 progress through the biogenesis pathway. Stable binding of Nog2 to H92 is a prerequisite for recruitment of the Rix1/Rea1 complex, which drives maturation of the central protuberance. Subsequent GTP hydrolysis by Nog2 promotes its release from the pre-60S^24^. Because Nog2 occludes the binding site for the export adapter protein Nmd3, once Nog2 is released, Nmd3 is free to bind to escort the pre-60S to the cytoplasm (Figure 4E).

In human cells, G4469 (the equivalent of G2922) is also methylated^25^. Because Nog2 (GNL2 in humans) is highly conserved between yeast and humans, we anticipate a similar functional and structural role for Gm4469 in assembly of the human large ribosomal subunit. Furthermore, surveillance of rRNA modification by orthologous GTPases may well extend beyond nuclear ribosome assembly in eukaryotes. Several studies examining the human mitochondrial ortholog of Nog2, MTG1 (GTPBP7), suggest this enzyme binds to the maturing H92 of the mitochondrial large subunit, and may directly sense modifications of universally conserved A-loop bases including Um3039 or Gm3040 ^26–28^. Furthermore, the Nog2 bacterial ortholog RbgA has been shown to interact with H92 in *B. subtilis* ribosome assembly^29^, where the A-loop base Gm2582 (analogous to *S. cerevisiae* Gm2922) is suggested to be 2’-O-methylated^30^. The structures of bacterial and mitochondrial Nog2 orthologs have conserved histidine residues thought to play a role in catalysis^31^, either by coordinating a water molecule for the first nucleophilic attack step of GTP hydrolysis, or controlling access to the gamma phosphate. Intriguingly, Gm2922 fills an analogous position in the active site channel of Nog2, potentially occluding access of water to the γ phosphate of GTP (Figure S6C). Alternatively, the 2-amino group of Gm2922 could play a direct role in coordinating water. Though these enzymes have experienced more than one billion years of evolutionary divergence, their common structural features and binding location on the nascent large ribosomal subunit suggests that RNA modification play a critical role in regulating their enzymatic functions. Taken together, our results suggest a new paradigm for RNA modifications in gating assembly or function of macromolecular machines in the cell.

## Acknowledgments

We thank A. Brillot for assistance with cryo-electron microscopy and data acquisition. We also thank I. Hoskins for assistance with designing primers for TileSeq and J. Weile for helpful discussion. We are grateful to B. Xhemalce and J. Huibregtse for helpful comments on the manuscript, and to all members of the Johnson, Taylor and Sarinay-Cenik labs for helpful discussions.

## Author contributions

Conceptualization: J.N.Y, A.W.J; Investigation: J.N.Y., J.P.K.B., J.J.B.; Formal analysis: J.N.Y., J.P.K.B.; Visualization: J.N.Y., J.P.K.B., J.J.B.; Writing, original draft: J.N.Y., D.W.T., A.W.J.; Writing, review and editing: All authors; Supervision: D.W.T., A.W.J.; Funding acquisition: D.W.T., A.W.J.

## Data and materials availability

The raw sequencing data are deposited in the Sequencing Read Archive (SRA) with accession number xxxxxxx. The structure of the Nog2-3xFLAG unmodified pre-60S subunit and its associated atomic coordinates have been deposited into the Electron Microscopy Data Bank (EMDB) and the Protein Data Bank (PDB) as EMDB-xxxx, and PDB code xxxx, respectively. All other data are available in the manuscript or supplementary materials. Requests for strains or plasmids will be fulfilled by the lead contact author, A.W.J., upon request.

## Supplemental information

### Methods

#### Strains, plasmids and cell growth

Strains are listed in Table 2, plasmids in Table 3 and oligonucleotides in Table 4. All yeast strains are derivatives of BY4741 and were grown at 30 °C. Golden Gate assemblies were performed using BsaI-HF or Esp3I enzymes from New England Biolabs. All site-directed mutagenesis was performed using inverse PCR or Gibson assembly with mutagenic oligonucleotides. All plasmid assemblies requiring a PCR-amplified insert were verified via Sanger sequencing. Yeast transformations were all conducted via the PEG lithium acetate method^32^. Yeast Toolkit (YTK) plasmids were a gift from J. Dueber (Addgene kit #1000000061)^33^.

**Table 2.** Yeast strains.

| Strain | Relevant Genotype | Source |
| --- | --- | --- |
| AJY4406 | *MATa ura3Δ0 leu2Δ0 his3Δ1*<br>*NatR::pGAL1:NOG2* | This study |
| AJY4425 | *MATa ura3Δ0 leu2Δ0 his3Δ1 KanMX::pGAL1:SPB1* | This study |
| AJY4427 | *MATa ura3Δ0 leu2Δ0 his3Δ1 KanMX::pGAL1:SPB1*<br>*NOG2Δintron-3xFLAG-6xHis* | This study |

**Table 3.** Plasmids.

| Plasmid | Relevant Sequence | Source |
| --- | --- | --- |
| pAJ4247 | *pPGK1:SpCas9 ptF(GAA)B:sgRNA::sfGFP URA3 CEN KanR* | <sup>35</sup> |
| pAJ4395 | *NOG2 ΔBsal Δintron LEU2 CEN AmpR* | This study |
| pAJ4810 | *pPGK1:SpCas9 pTF(GAA)B:sgRNA-NOG2 URA3 CEN KanR* | This study |
| pAJ4817 | *NOG2 ΔBsal Δintron S208A LEU2 CEN AmpR* | This study |
| pAJ4819 | *NOG2 ΔBsal Δintron R389S LEU2 CEN AmpR* | This study |
| pAJ4821 | *NOG2 ΔBsal Δintron S208A R389S LEU2 CEN AmpR* | This study |
| pAJ4841 | *SPB1 HIS3 CEN AmpR* | This study |
| pAJ4842 | *spb1-D52A HIS3 CEN AmpR* | This study |
| pAJ4852 | *SRG2922 LEU2 2μ AmpR* | This study |
| pAJ4876 | *NOG2 ΔBsal Δintron LEU2 2μ AmpR* | This study |
| pAJ4881 | *NOG2-H392R ΔBsal Δintron LEU2 2μ AmpR* | This study |
| pAJ4893 | *NOG2-T195R ΔBsal Δintron LEU2 2μ AmpR* | This study |
| pAJ4890 | *RPL25-eGFP NOP56-mRFP URA3 CEN AmpR* | This study |
| pAJ5103 | *CONS sfGFP-dropout CON1 LEU2 CEN AmpR* | This study |
| pAJ5104 | *CONS sfGFP-dropout CON1 HIS3 CEN AmpR* | This study |
| pAJ5125 | *LEU2 CEN AmpR* | This study |
| pAJ5126 | *HIS3 CEN AmpR* | This study |
| pAJ5136 | *LEU2 2μ AmpR* | This study |

**Table 4.**
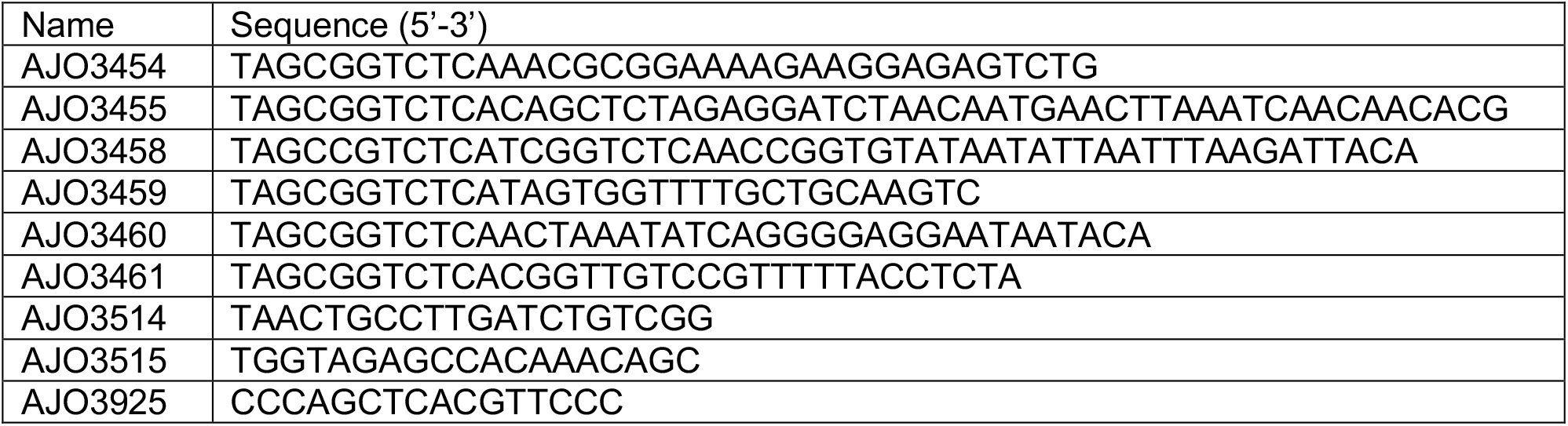
Primers.

Strain AJY4406 (*NatR::pGAL1:3xHA-NOG2*) was generated via PCR amplification of a modular *NatR:pGAL1:3xHA* cassette with oligonucleotides containing *NOG2* promoter and coding sequence, as previously described^34^. Strain AJY4425 (*KanMX::pGAL1:SBP1*) was constructed similarly. Strain AJY4427 (*KanMX::pGAL1:SPB1 NOG2Δintron-3xFLAG-6xHis*) was generated via a YTK-compatible Cas9 protocol described previously^35^. A Cas9 sgRNA (5’-TTTGATTCTTTCCAACCAAG-3’) was designed to target the intron within *NOG2* and assembled into sgRNA dropout vector pAJ4247 to construct the constitutive sgRNA plasmid pAJ4810; strain AJY4425 was then co-transformed with pAJ4810 and a linearized repair template corresponding to *NOG2* without intron and containing a C-terminal 3xFLAG-6xHis tag (derived from digestion of pAJ4853 with EagI). Candidate *NOG2 Δintron-3xFLAG* recombinants were screened for loss of the *NOG2* intron by colony PCR and for Nog2-3xFLAG expression by Western blot. A successful recombinant was finally streaked on yeast peptone galactose (YP-Gal) media, and colonies screened for URA3 auxotrophy to verify loss of pAJ4810.

#### *PCR mutagenesis of* NOG2

Plasmid pAJ4395 was constructed by amplifying the *NOG2* locus (∼500 nt upstream of the *NOG2* start codon to ∼80 nt downstream of the *NOG2* stop codon) from genomic DNA purified from yeast strain BY4741, and cloning it into vector pAJ5103 (*CONS-sfGFP dropout-CON1 LEU2 CEN AmpR*) assembled from YTK parts^33^. The internal BsaI site in the *NOG2* coding sequence was replaced with a synonymous coding sequence (G444>A), and the single intron within the *NOG2* locus was removed. The *NOG2* open reading frame was mutagenized using Taq polymerase (New England Biolabs), with amplifying primers AJO3460 and AJO3461, containing BsaI sites for Golden Gate assembly. To alleviate bias in the type and rate of mutations, dNTP concentrations were adjusted to 0.2 mM dATP and dGTP, and 1 mM dTTP and dCTP^36^. After 5 amplification cycles, manganese chloride was added to a final concentration of 500 µM^37^. PCR was then continued for another 20 cycles. The promoter (∼500 nt) and terminator (∼200 nt) regions of *NOG2* were amplified separately from BY4741 genomic DNA, with a high-fidelity polymerase, using oligonucleotides AJO3454 x AJO3459 (*NOG2* promoter) and AJO3458 x AJO 3455 (*NOG2* terminator), all of which contained compatible 5’ BsaI sites for directional assembly.

Mutagenized *NOG2* was combined with the PCR products encoding its promoter and terminator regions in a BsaI Golden Gate assembly reaction, and with GFP dropout vector pAJ5103. The assembled product was used to transform competent DH5α *E. coli* (New England Biolabs), and transformants (∼10^5^) were selected on ampicillin plates. Fewer than 1 in 10^4^ colonies expressed GFP, indicating successful library assembly. Several colonies were sequenced to verify the presence, quantity and uniformity of mutations, and the rest were pooled and bulk DNA prepared via Midiprep (Qiagen).

The mutagenized library prep was digested with BsaI to eliminate any remaining vector background and used to transform strain AJY4406. Transformants (∼5 × 10^4^) were selected on synthetic solid medium containing galactose and lacking leucine, pooled, and frozen in a glycerol stock. Cells from this stock were plated onto synthetic medium containing glucose and lacking leucine, thereby selecting transformants that could complement repression of the endogenous *NOG2* locus. Colonies viable on glucose (∼5 × 10^4^) were pooled and frozen in a glycerol stock.

To estimate sequencer errors, a wild-type control was carried out in which strain AJY4406 was transformed with plasmid pAJ4395, harboring the unmutagenized *NOG2* coding sequence. These transformants were grown in synthetic liquid media lacking leucine, with galactose or glucose as a carbon source for the input and selection controls, respectively.

#### Sequencing and analysis of mutagenized yeast libraries

Plasmid extraction and all subsequent steps were performed in duplicate. Briefly, plasmid DNA was extracted from ∼3 × 10^9^ cells and amplified with vector-specific oligonucleotides AJO3514 and AJO3515. These amplicons were subsequently PCR amplified using 15 interleaved primer pairs, each yielding an amplicon of 150 base pairs, flanked by sequence for Illumina adapter priming. Amplicons for sequencing were pooled according to sample, and multiplexing Illumina adapters were attached in a final PCR step. The pools were sequenced in a single run on an Illumina NextSeq platform, using 150 bp paired-end reads. Reads were aligned to the *NOG2* open reading frame using BowTie2^38^ and split into their corresponding tiles with a custom Python script. Mutations were counted and normalized using the TileSeq Analysis package (Version 1.5)^8^. These mutational counts were analyzed and the complete phenotypic profile imputed and displayed using the POPCode Analysis Pipeline^8^. Custom Python scripts were used to calculate median fitness scores from the complete analysis, and map these scores onto the atomic model of Nog2.

#### Sucrose gradient sedimentation

Strain AJY4425 was co-transformed with the following paired combinations of plasmids: pAJ4841/pAJ5136, pAJ4842/pAJ5136, pAJ4842/pAJ4852, and pAJ4842/pAJ4881. The transformants were inoculated into synthetic media lacking leucine and histidine and containing galactose, and grown overnight at 30°C to saturation. The transformants were then diluted 1:200 into synthetic media lacking leucine and histidine and grown overnight at 30 °C to repress the expression of genomic *SPB1*. Cells from these cultures were finally diluted into the same synthetic media, grown to mid-log phase (OD_600_ ≈ 0.6,) then treated with 100 µg/mL cycloheximide and shaken for 30 minutes at 30°C. Treated cells were centrifuged and stored at -80°C. Thawed cells were washed and resuspended in lysis buffer (20 mM HEPES-KCl pH 7.4, 100 mM KCl, 2 mM MgCl_2_, 5mM β-mercaptoethanol, 100 µg/mL cycloheximide, 1 mM each of PMSF and benzamidine, and 1 μM each of leupeptin and pepstatin). Extracts were prepared by glass bead lysis (two 30-second rounds of vortexing) and clarified by centrifuging for 15 minutes at 18,000 g at 4°C. 4.5 A_260_ units of clarified extract were loaded onto a 7-47% sucrose gradient made in the same buffer lacking the protease inhibitors. Gradients were centrifuged for 2.5 hours at 40,000 rpm in a Beckman SW40 rotor, and RNA content monitored at 254 nm using an ISCO Model 640 fractionator.

#### RNAse T1 digestion and primer extension

Strain AJY4425 was co-transformed with the following paired combinations of plasmids: pAJ4841/pAJ5136, pAJ4842/pAJ5136, pAJ4842/pAJ4852, pAJ4842/pAJ4881. The transformants were inoculated into synthetic media lacking leucine and histidine and containing galactose, and grown overnight at 30°C to saturation. To repress the expression of genomic *PGAL1-SPB1*, the cells were diluted 1:200 into pre-warmed media containing glucose and grown overnight at 30°C to saturation. Finally, to dilute mature ribosomes synthesized prior to Spb1 depletion, the cells were again diluted 1:2000 into fresh, pre-warmed SD-Leu/His media containing glucose and cultured at 30°C until early exponential phase (OD_600_ ≈ 0.3). Cells were harvested and stored at -80°C. Total RNA was prepared using the acid-phenol-chloroform method as previously described^39^. To assay the methylation status of G2922, equivalent amounts of 25S rRNA were dried by speed vacuuming and resuspended in annealing buffer (40 mM PIPES pH 7.0, 400 mM NaCl, 1 mM EDTA). A mixture of RNA and P^32^-labeled AJO3925 (5’-CCCAGCTCACGTTCCC-3’) was heated at 95°C for 3 minutes and cooled to 4°C using a ramp-down rate of 10 seconds per degree centigrade to promote annealing in a final volume of 25 µL. After annealing, 2 units of RNase T1 (Ambion) were added to 10 µL of the annealing reaction and incubated at 25°C for 60 minutes. Reverse transcription reactions using MMLV reverse transcriptase (Invitrogen) were performed according to the manufacturer’s instructions, using 4 µL of each RNase T1 digest reaction as template in a final reaction volume of 10 µL. Reactions were carried out at 37°C for 50 minutes followed by inactivation at 70°C for 15 minutes. 1 volume (10 µL) of 2X TBE-urea sample buffer (89 mM Tris, 89 mM Boric acid, 2 mM EDTA pH 8.0, 7M urea, 12% Ficoll, 0.01% bromophenol blue, 0.02% xylene cyanol FF) was added to each sample and heated at 70°C for 15 minutes. 2 µL of each sample was separated on a 12% TBE-urea gel, dried on filter paper, and exposed to a phosphoscreen overnight. Signal was detected by phosphorimaging on a GE Typhoon FLA9500.

#### Fluorescence microscopy

Strain AJY4425 harboring pAJ4890 was co-transformed with the following paired combinations of plasmids: pAJ4841/pAJ5136, pAJ4842/pAJ5136, pAJ4842/pAJ4852, pAJ4842/pAJ4872, pAJ4842/pAJ4881. Strains were grown to saturation in synthetic media containing galactose and lacking uracil, leucine and histidine, then subcultured for 16 hours in media containing glucose to deplete endogenous Spb1. Cells were then freshly diluted 1:20 and grown for 4 hours in media containing glucose. Cells were fixed for 30 minutes with 3.7% freshly prepared formalin, washed three times with cold 100 mM potassium phosphate (pH 6.4), and incubated for 5 minutes at room temperature with 0.1% Triton X-100. DAPI was added to 2 µg/mL and cells were incubated for another 3 minutes at room temperature, then washed three times with cold PBS (pH 7.4) and stored at 4 °C. Fluorescence and brightfield micrographs were recorded on a Nikon E800 microscope fitted with a 100X Plan Apo objective and a Photometrics CoolSNAP ES camera controlled by NIS-Elements software.

#### Immunopurification

Strain AJY4427 (*KanMX::pGAL1:SPB1 NOG2-3xFLAG-6xHis*) was transformed with wild-type SPB1 (pAJ4841) or *spb1-D52A* (pAJ4842) and grown for 24 hours in media lacking histidine and containing galactose, followed by 24 hours growth in media containing glucose to deplete endogenous Spb1. Cells were then diluted in the same medium and grown to mid log-phase (OD_600_ of 0.6). For analysis by PAGE and MS, cells were harvested and frozen at -80 °C. All subsequent steps were performed at 0-4°C. Cell pellets were thawed and resuspended in IP binding buffer (50 mM Tris pH 7.6, 100 mM NaCl, 5 mM MgCl_2_, 0.05% NP-40, 0.5 mM TCEP) and broken with glass beads. Cell lysate was clarified by two subsequent centrifugation steps of 5 minutes at 3,000 g followed by 20 minutes at 18,000 g, and the lysate was incubated with αFLAG-conjugated agarose beads (Millipore-Sigma) for 2 hours with rotation at 4 °C. Lysate was removed and beads were washed three times with IP wash buffer (50 mM Tris pH 7.6, 100 mM NaCl, 5 mM MgCl_2_, 0.01% octyl-β-glucoside, 0.5 mM TCEP), after which the Nog2-3xFLAG complex was eluted for 90 minutes with IP wash buffer containing 350 µg/mL 3xFLAG peptide. For analysis by cryo-EM, cells were flash frozen as a 75% cell slurry in IP buffer, and broken under cryogenic conditions in a Retsch mixer mill with 6 cycles of 3 minutes at 15 Hz. Subsequent steps were as described for analysis by PAGE and MS above. Eluates were either frozen and stored for PAGE and MS analysis, or else directly applied to cryo-electron microscopy grids.

#### Mass spectrometry

Protein identification was provided by the University of Texas at Austin Center for Biomedical Research Support Proteomics Facility (RRID:SCR_021728). Samples were excised from SDS-PAGE gels, trypsin digested, desalted and run on a Dionex LC and Orbitrap Fusion 2 for LC-MS/MS for 30 minutes (single band) or 120 minutes (whole-lane), and analyzed using PD 2.2 and Scaffold 5. For semi-quantiative analysis, peptide spectral counts were normalized to protein molecular weight, and P-values for enrichment were calculated from a paired *t*-test using the Cyber-T web server^40^ without Bayesian correction.

#### Cryo-electron microscopy

Quantifoil R1.2/1.3 grids coated with an ultrathin layer of amorphous carbon (Electron Microscopy Sciences) were glow-discharged for 1 minute at 25 mA. Using a Mark IV VitroBot (FEI), sample was applied to freshly glow-discharged grids at 4 °C and 100% humidity, immediately blotted for 2 seconds at a force setting of 0, then plunged into liquid ethane and stored under liquid nitrogen until data collection. Microscopy data were collected in two separate rounds at the University of Texas at Austin Sauer Structural Biology Center on a Titan Krios microscope (FEI) operating at 300 kV, equipped with a K3 Summit direct electron detector (Gatan). To alleviate particle orientation bias observed in the first collection, 6,768 movies were recorded at 0° tilt and 6,732 movies were recorded at 30° tilt. In the second round of data collection, 12,456 movies were collected at 0° tilt, for a total of 25,956 movies. Movies were acquired using SerialEM^41^ and recorded as 20 frames over 4 seconds for a total electron dosage of ∼70 e^-^ per Å. Movies were recorded at a nominal magnification of 22,500x and pixel size of 0.81 Å. Using cryoSPARC Live for on-the-fly processing, movies were motion-corrected and dose-weighted, CTF estimates performed, and a total of 293,350 particles were picked using a 60S template. Following on-the-fly 2D classification to separate clean (pre-)60S projections, particles were exported to cryoSPARC^42^ (version 3.2), where multiple subsequent rounds of 2D classification resulted in a total of 194,497 particles. These particles were used for *ab initio* reconstruction and 3D heterogeneous refinement to separate a total of 120,722 nucl(eol)ar particles with at least partial Nog2 occupancy, which were subjected to consensus non-uniform refinement. Using the csparc2star.py function in pyEM^43^, these particles with were exported to RELION^44^ (version 3.1.3), where a 3D classification scheme using a soft mask around Nog2/H92 was used to further separate 86,273 Nog2-bound particles. These particles were re-imported into cryoSPARC for a 3D classification scheme with a soft mask around Rsa4, which separated 15,954 particles with Nog2 at high resolution. The final map was prepared using non-uniform refinement implemented in cryoSPARC, where FSC calculations and local resolution analysis were also carried out after final refinement.

#### Modeling, refinement and graphics

PDB 3JCT^10^ was first rigid-body docked into the final, unsharpened map, using UCSF ChimeraX^45^. Chains for uL11, Alb1, and YBL028C were added from PDB 6YLH^12^. The ribosomal RNA of H92 was fit by hand into the corresponding map density using COOT^46^. The model was relaxed using flexible molecular dynamics fitting with ISOLDE^47^, then finally subjected to real-space refinement as implemented in PHENIX^48^. All molecular graphics were prepared with UCSF ChimeraX^45^.

## Supplemental figure legends

**Figure S1.**
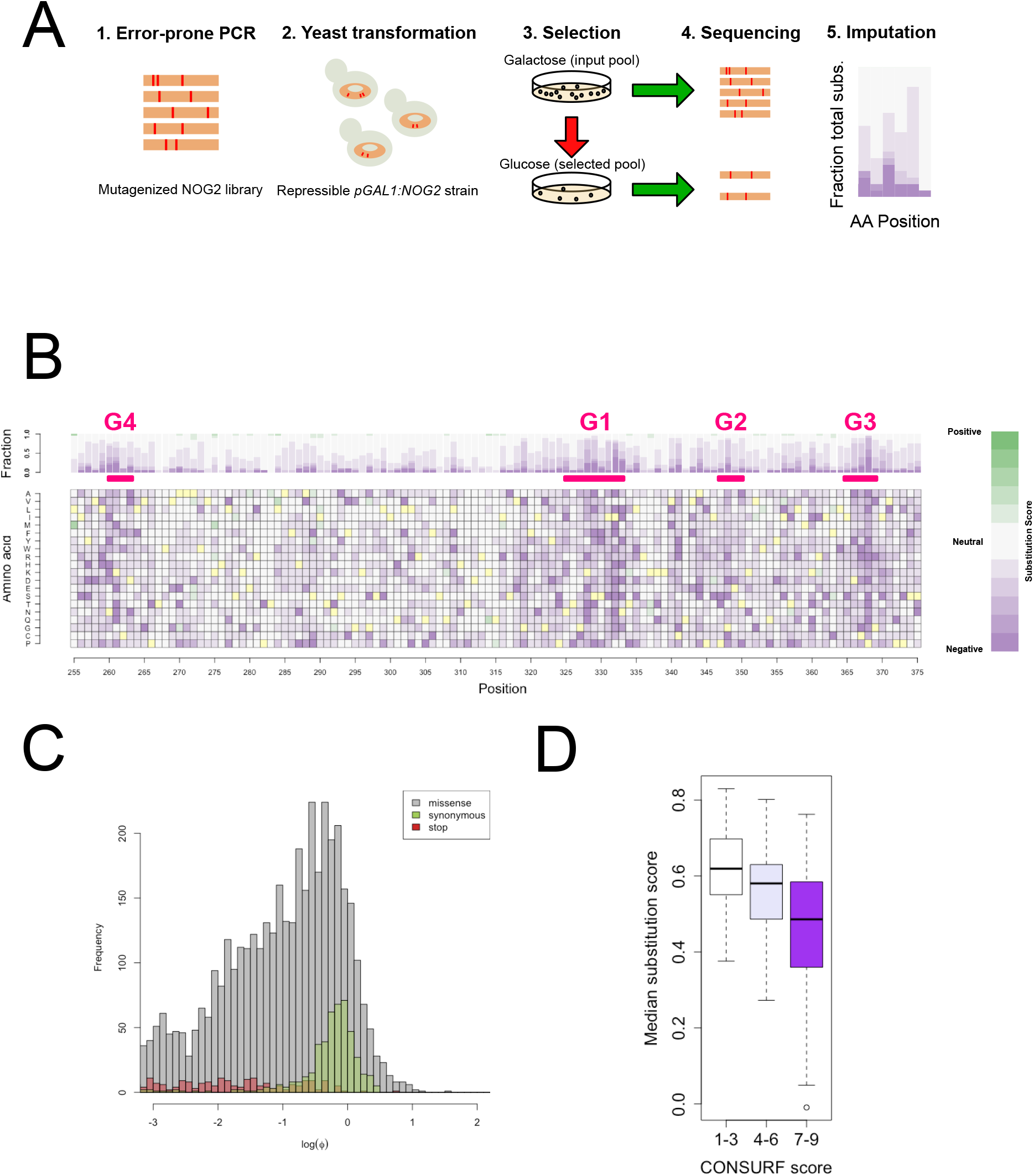
**A**. Scheme for high throughput mutagenesis and analysis of functional variants of *NOG2*. **B**. Complete fitness landscape for variants in the core GTP-binding motifs of *NOG2* (G4, G1, G2, G3; amino acids 260-369). **C**. Distributions of *observed* fitness scores (Φ) for nonsense (red), missense (gray) and synonymous (green) mutations. **D**. Correlation between per-residue median fitness score and ConSURF^49^ score (evolutionary conservation: scores 1-3, low; scores 4-6, medium; scores 7-9, high conservation).

**Figure S2.**
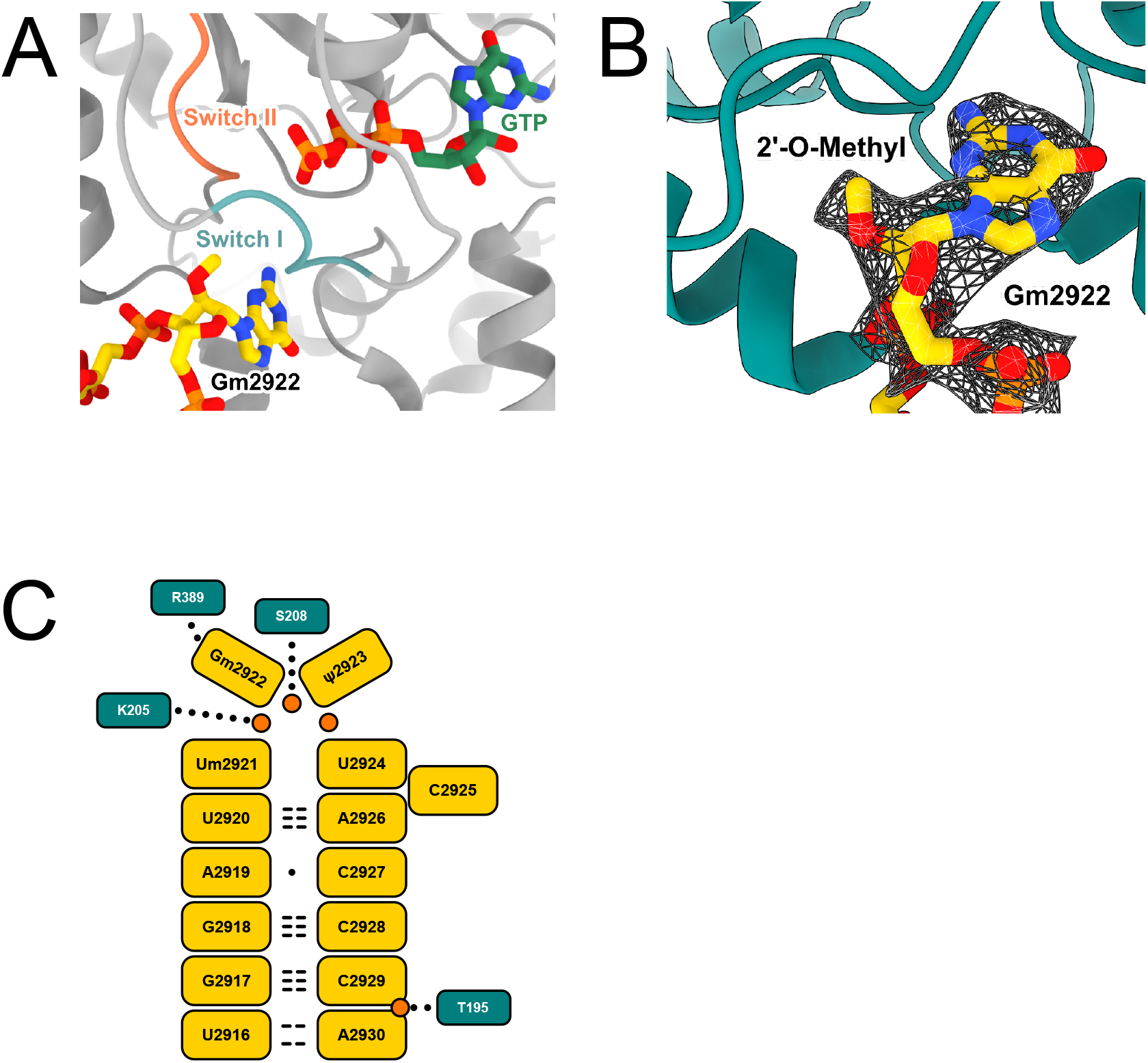
**A**. Modeling of active site of Nog2 with Gm2922 shown and Switch I (G2) and Switch II (G3) colored. **B**. Cryo-EM map of Nog2 (EMDB-6615) showing clear density for 2’-O-methyl group at G2922. **C**. Two-dimensional map of interactions between Nog2 amino acids and RNA elements of H92. Nucleic and amino acids are depicted as rectangles, relevant phosphates are shown as orange circles. Hydrogen bonds between bases are shown as dashed lines, and the wobble base pair between A2919 and C2927 is shown as a dot. Interactions between amino acids and H92 are depicted as dotted lines. Threonine 195 (T195) was selected as a candidate for a suppressing amino acid substitution (Figure 3).

**Figure S3.**
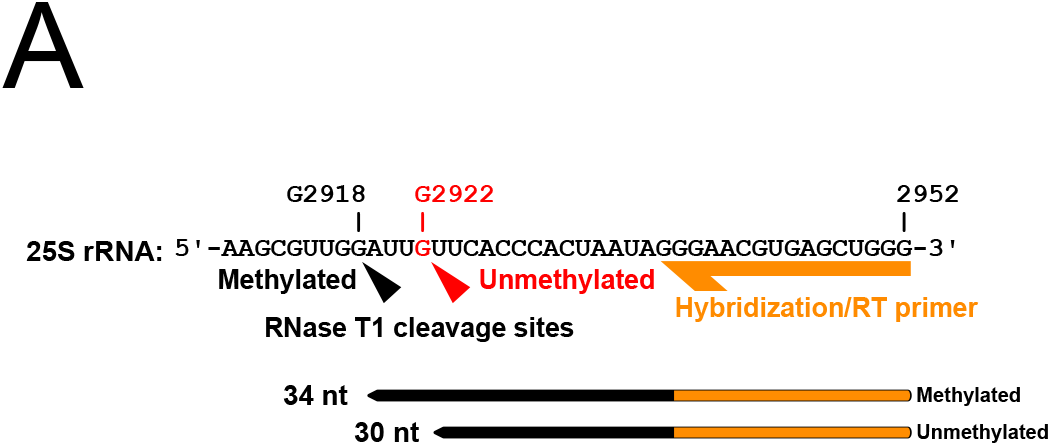
**A**. Schematic of primer extension probing 2’-O-methylation at G2922. RNase T1 cleaves 3’ to guanosines in single-stranded RNA. However, 2’-O-methylated guanosine is resistant to cleavage. Guanosines base-paired with the reverse transcription primer are also protected. After cleavage with RNase T1, reverse transcription will generate a 30 nt product if G2922 is sensitive to cleavage (i.e. not methylated), or a 34 nt product if G2922 is protected from cleavage (i.e. methylated).

**Figure S4.**
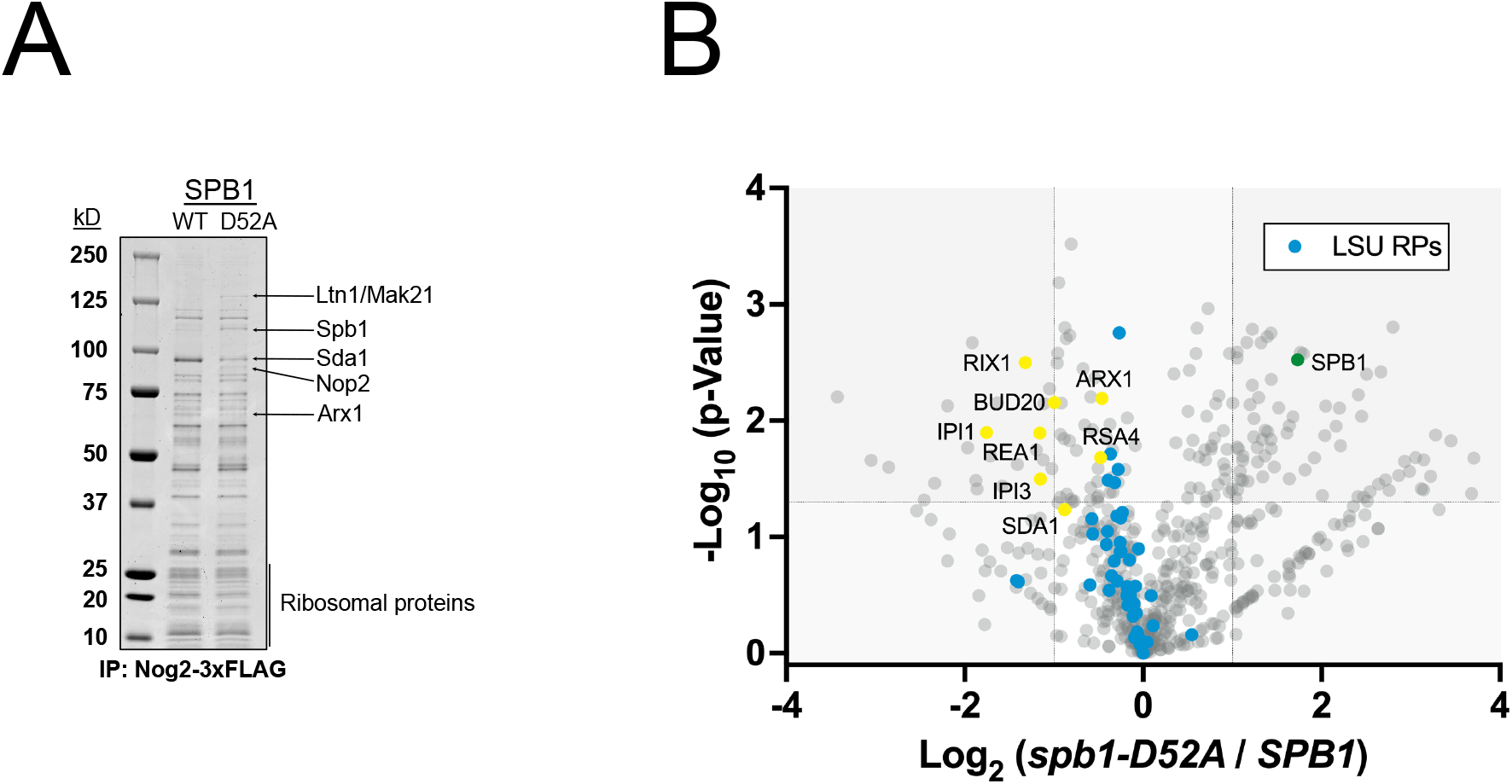
**A**. Nog2-3xFLAG immunopurifications from the indicated strains, stained with Coomassie blue. Proteins identified by mass spectrometry are indicated. **B**. Volcano plot showing log_2_ fold change of factors that coimmunopurify with Nog2-3xFLAG, in unmodified G2922 (*spb1-D52A*) versus modified Gm2922 (WT *SPB1*). Dashed lines intersecting x-axis indicate a log_2_ fold change of 1, and intersecting y-axis indicate a -log_10_ (*p*) value of 1.3 (*p* = 0.05).

**Figure S5.**
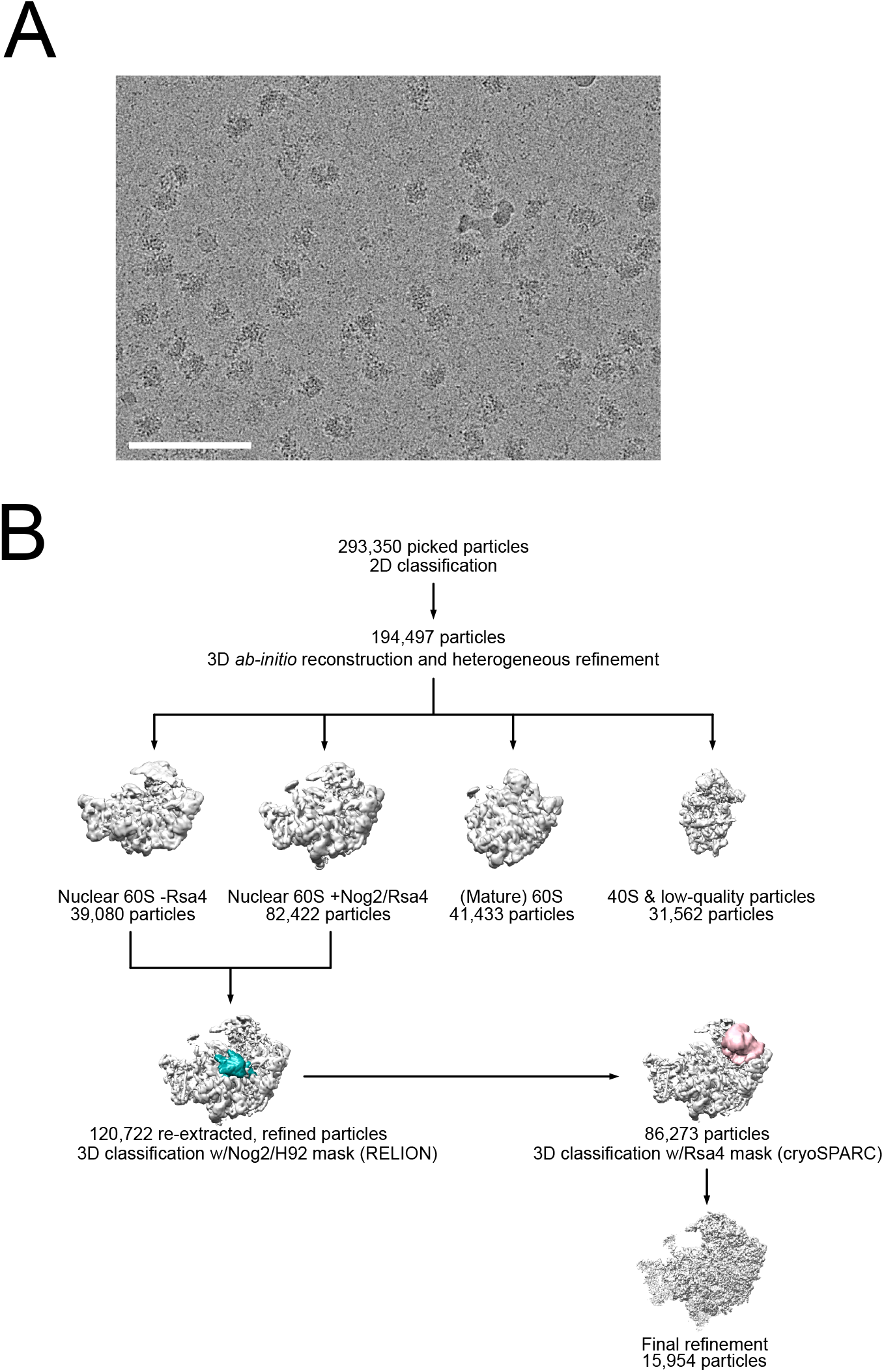
**A**. Representative micrograph of Nog2-3xFLAG particles immunopurified from the *pGAL:SPB1* strain AJY4425 expressing *spb1-D52A*. Scale bar = 100 nm. **B**. Cryo-EM data processing scheme by which the Nog2-3xFLAG structure was obtained.

**Figure S6.**
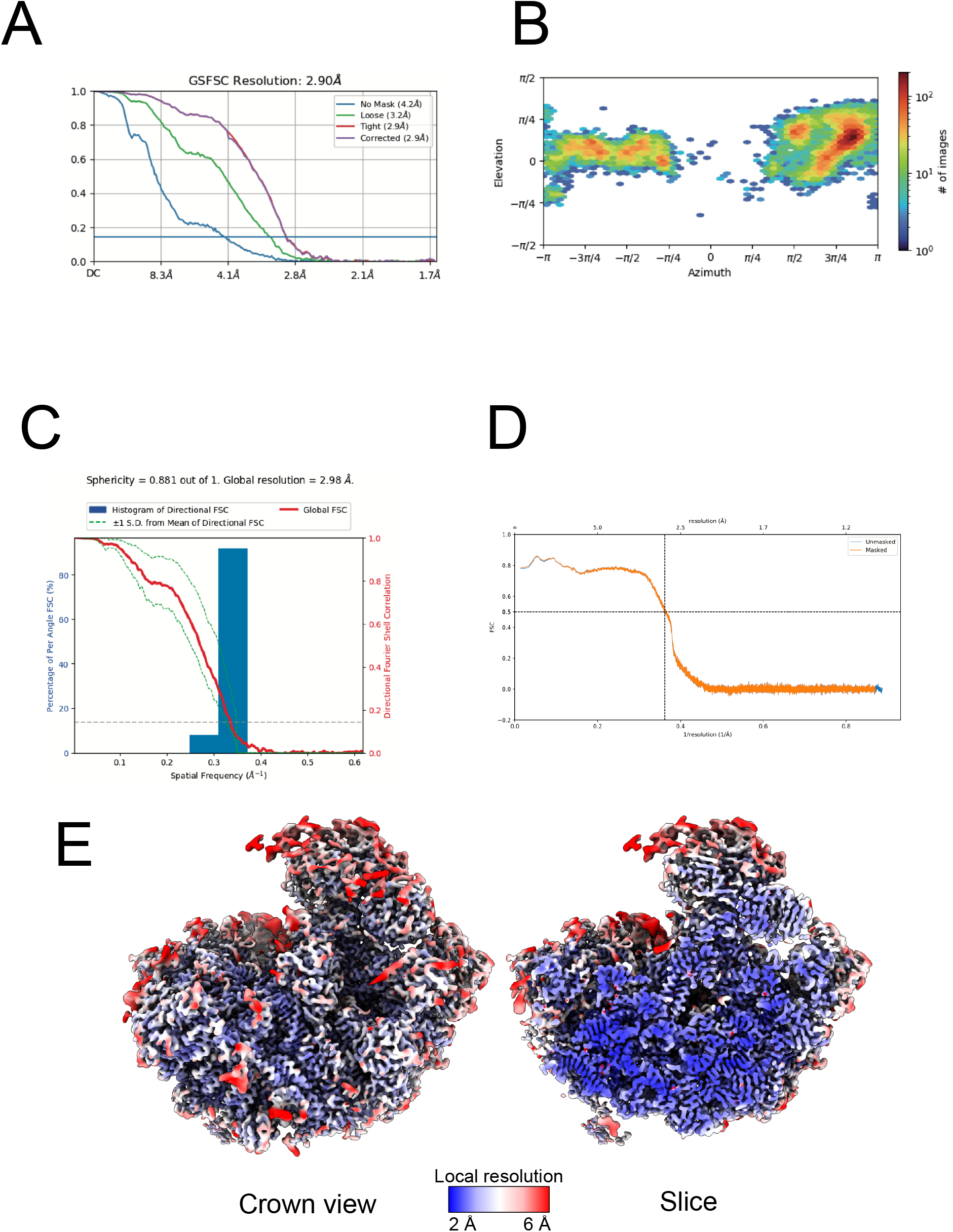
**A-B**. Fourier shell correlation (FSC) curves **(A)**, and distribution of viewing orientations **(B)** for the unmodified G2922 Nog2-3xFLAG structure. **C-D**. Directional **(C)** and map-to-model **(D)** FSC curves for the unmodified G2922 Nog2-3xFLAG structure. **(E)** Local resolution analysis of the overall unmodified G2922 Nog2-3xFLAG structure.

**Figure S7.**
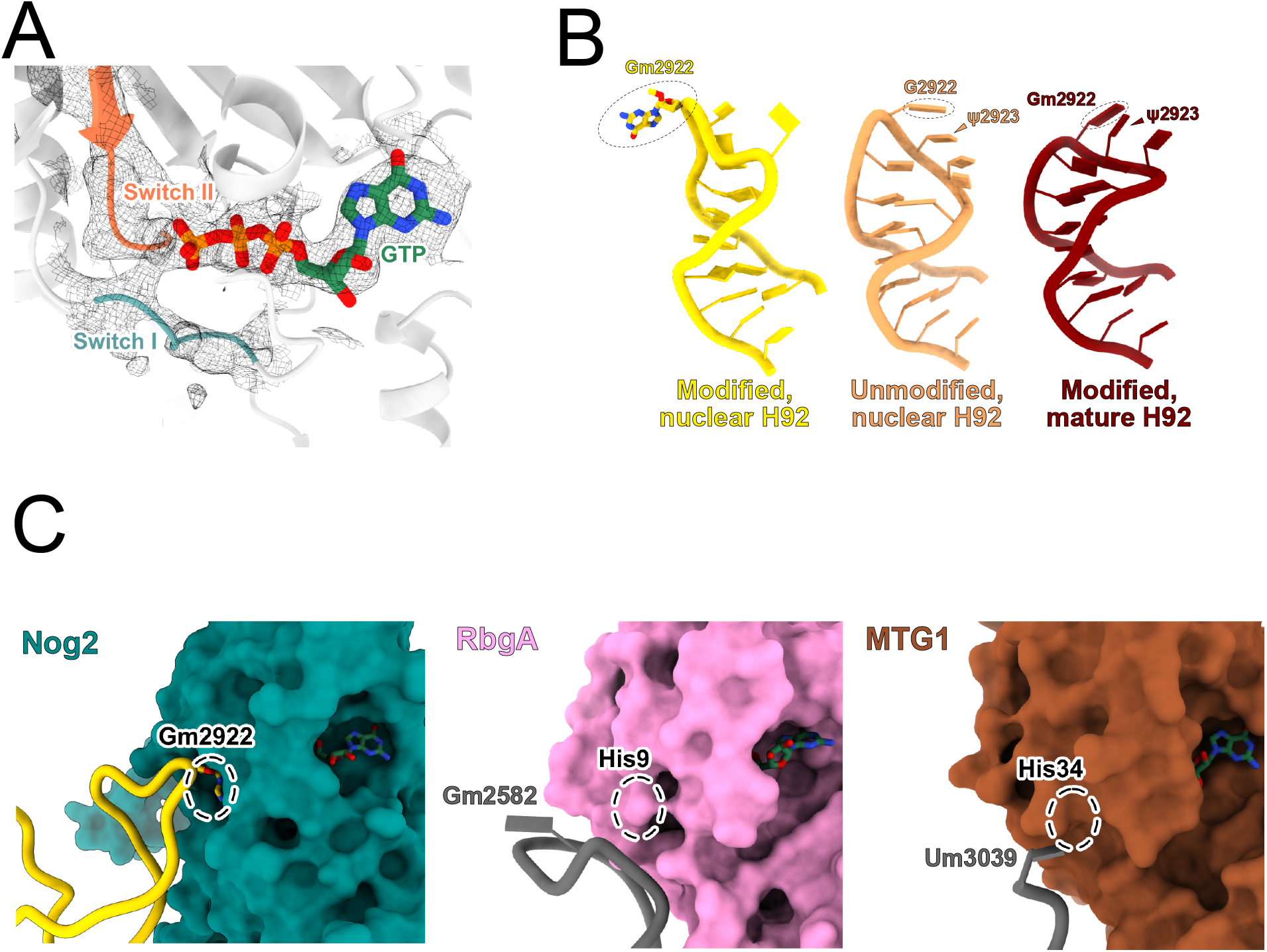
**A**. Active site of Nog2 in complex with pre-60S lacking G2922 methylation. Densities for GTP and ordered regions Switch I and Switch II are clearly observed. **B**. Cartoon comparisons between H92 with a modified Gm2922 engaged with Nog2 (left, PDB 3JCT), an unmodified G2922 (middle, this work) and the mature H92 from the yeast ribosome crystal structure (right, PDB 4V88). **C**. Comparison between the active sites of Nog2 (left, PDB 3JCT), bacterial ribosome biogenesis GTPase RbgA (middle, PDB 6PPK) and human mitoribosome biogenesis GTPase MTG1 (right, PDB 7PD3). The position of Gm2922, or locations of histidine residues implicated in catalysis, are indicated by dashed circles.

